# The Impact of Coronary Artery Disease on Brain Vascular and Metabolic Health: Links to Cognitive Function

**DOI:** 10.1101/2025.06.02.657532

**Authors:** Safa Sanami, Stefanie A. Tremblay, Ali Rezaei, Zacharie Potvin-Jutras, Dalia Sabra, Brittany Intzandt, Christine Gagnon, Amélie Mainville-Berthiaume, Lindsay Wright, Mathieu Gayda, Josep Iglesies-Grau, Anil Nigam, Louis Bherer, Claudine J. Gauthier

## Abstract

**Background:** Coronary artery disease (CAD) is the leading cause of mortality worldwide and is associated with an increased incidence of cognitive decline, however the pathological mechanisms linking CAD to brain and cognitive health remain unclear. Prior research has identified regional deficits in cerebral blood flow (CBF) and cerebrovascular reactivity (CVR), a measure of vascular reserve, in patients with CAD. However, the impact of these cerebrovascular deficits on cognition has not been explored, nor has the effect of CAD on cerebral metabolic health. This study aims to fill these gaps by investigating how CAD influences cerebral vascular and metabolic health, and how these alterations relate to cognitive function across multiple domains.

**Methods:** Quantitative magnetic resonance imaging (MRI) was employed to measure a comprehensive profile of cerebral vascular and metabolic health, including CBF, CVR, cerebral metabolic rate of oxygen (CMRO₂), and oxygen extraction fraction (OEF). Cognition was assessed using a validated neuropsychological battery from which composite scores were extracted, reflecting executive functions, working memory, processing speed, and verbal episodic memory.

**Results:** A total of 35 patients with CAD and 37 healthy controls were included in the final analysis. Patients with CAD demonstrated widespread impairments in both cerebral vascular and metabolic health, characterized by lower CBF, CVR, and CMRO₂, and increased OEF, indicative of insufficient oxygen delivery. Notably, lower CVR was associated with poorer executive function, suggesting a specific role of vascular reserve for preserving executive functions. Furthermore, higher OEF was associated with poorer working memory, showing the importance of preserved oxygen consumption for maintaining cognitive function.

**Conclusion:** This study reveals that CAD is associated with impaired cerebral vascular and metabolic health, providing a pathological basis for cognitive decline. Specifically, reduced CVR and elevated OEF emerged as sensitive biomarkers linking impaired brain health and cognition. These markers hold promise for guiding future interventions aimed at preserving cognitive health in patients with CAD.

**Clinical Perspective:** *What Is New?:*

- Coronary artery disease is associated with an increased incidence of cognitive decline but the biological mechanisms underlying these changes in cognition are unclear.
- This is the first study to comprehensively demonstrate profound and widespread deficits in both cerebral vascular and oxidative metabolic health in patients with coronary artery disease.
- Cerebrovascular reactivity and oxygen extraction fraction may be sensitive biomarkers of early cognitive decline in coronary artery disease.

*What Are the Clinical Implications?:*

- The widespread impairments in cerebral vascular and metabolic health suggest that coronary artery disease has profound consequences on the brain.
- Poor vascular reserve and an imbalance between oxygen delivery and usage may be crucial determinants of cognitive health. These results highlight the limitations of traditional biomarkers in understanding the brain’s vulnerability and the need for these novel and quantitative biomarkers of cerebral physiological health.
- These findings suggest that cerebral vascular and metabolic deficits exist even in the absence of overt cognitive decline in coronary artery disease, emphasizing the need for early detection and preventative strategies.

## Introduction

Coronary artery disease (CAD) is the leading cause of death worldwide and the most common chronic heart condition, affecting over 300 million people ^1^. With improved treatments prolonging survival, the downstream effects of CAD on the brain are now emerging, including an increased risk of cognitive decline, dementia, and stroke ^2^. Because these diseases take decades to develop, gradual and profound impacts on brain health likely start accumulating long before the onset of neurological disease in CAD patients, but these early changes are not well documented and cannot therefore be reversed or prevented.

Individuals with CAD have a 45% higher likelihood of developing cognitive impairment or dementia ^3^. Commonly affected domains include global cognition, attention, memory, processing speed, and executive function ^4–6^. The brain changes that underlie these cognitive deficits are poorly understood however. Structural brain alterations, including reduced gray matter volume, cortical thinning, and white matter lesions, have been reported in CAD ^7,8^. While these changes are associated with cognitive decline ^9^, they represent entrenched pathology that is difficult to reverse. In contrast, physiological dysfunctions — such as impairments in cerebral vascular and metabolic function — occur earlier and are more easily reversible ^10–12^.

Growing evidence suggests that CAD negatively impacts cerebrovascular function. Previous studies have observed regional reductions in both cerebral blood flow (CBF) ^10^ and cerebrovascular reactivity (CVR) in CAD ^10,13^. CVR represents the increase in CBF that occurs in response to a vasodilatory challenge such as elevated CO_2_ and is considered a measure of vascular reserve. Notably, CBF can be partially restored through cardiac rehabilitation ^10,12^, highlighting cerebrovascular function as an early and modifiable marker of brain health in CAD. The link between cerebrovascular health—specifically CBF and CVR—and cognitive function, while understudied in CAD, has been documented in various neurological conditions. Higher CBF has been consistently linked to improved cognitive performance across several domains in Alzheimer’s disease ^14^, mild cognitive impairment ^15^ and in healthy individuals ^16^. Though less studied, reduced CVR consistently relates to cognitive deficits, especially in executive functions and memory, in Alzheimer’s disease, vascular cognitive impairment, and healthy aging ^17–19^, as well as in one study in CAD using ultrasound ^13^. Given the consistent associations found across other clinical conditions, it is highly plausible that similar cerebrovascular deficits in CAD contribute to lower cognitive performance.

Cerebrovascular health is crucial for adequate substrate delivery, yet the brain’s capacity to function also relies on preserved metabolic processes. The human brain meets its substantial energy demands primarily through glucose utilization and oxidative metabolism ^20^. Though largely unstudied, there are emerging indications that metabolism and mitochondrial function may be affected in CAD ^21^. Should oxygen usage be compromised, this disruption could contribute to the cognitive deficits often reported in CAD. Metabolic deficits and mitochondrial dysfunction may lead to lower cerebral metabolic rate of oxygen consumption (CMRO₂) ^22,23^, as observed in individuals with Alzheimer’s and Huntington’s disease ^22,24^ and to a lesser extent in aging ^25^. An imbalance between delivery and consumption of O_2_, measured as high oxygen extraction fraction (OEF), a parameter reflecting the proportion of oxygen extracted from blood by brain tissue expressed as a percentage, is also likely present, as observed in individuals with high vascular risk factor load, with and without cognitive impairment ^26,27^. This high OEF would have clinical relevance given that it could be indicative of tissue at risk for ischemic damage ^28^. However, the presence of these types of metabolic deficits in CAD is currently unknown, as is their potential link with cognition.

The present study addresses these gaps by exploring the associations between cerebral vascular and metabolic deficits and cognition in CAD patients using quantitative MRI. We hypothesize that: 1. CAD patients will exhibit poorer cerebrovascular health, observed as reduced CBF and CVR compared to healthy controls (HC); 2. CAD patients will show reduced oxidative metabolism and an imbalance between oxygen usage and delivery, observed as decreased CMRO₂ and increased OEF compared to HCs; and 3. Better cerebral vascular and metabolic health will be associated with better cognitive performance. By comprehensively exploring brain physiological changes in CAD, this study will provide important insight into the long-term brain changes caused by CAD and pave the way for earlier detection and targeted interventions.

## Methods

### Participants

Ninety-nine participants of 50 years and above were recruited, from which 87 completed the study (46 CAD patients and 41 HCs, matched for age and sex). Out of 12 participants that discontinued their participation, five participants did not complete the study due to interruptions during the COVID-19 pandemic, two participants were excluded due to claustrophobia, two participants were excluded due to discomfort during the MRI and three participants chose not to participate for personal reasons (e.g. loss of interest, issues with scheduling). Of the 87 remaining participants, twelve had only 5 minutes of resting CBF data instead of the full acquisition including hypercapnia due to discomfort during hypercapnia and these were included in CBF analysis. Only 75 participants had a complete dataset. To ensure sufficient data quality, we excluded three participants with low-quality data based on the following criteria: pseudo-continuous arterial spin labeling (pCASL) temporal signal-to-noise ratio (tSNR) below 0.5 and excessive motion during the acquisition, defined as translational movements greater than 2mm, for a final sample of 35 CAD patients with CAD and 37 control subjects with hypercapnia (Table 1 shows demographics for participants with respiratory data; Table S1 includes all participants). The study received approval from the Comité d’éthique de la recherche et du développement des nouvelles technologies (CÉRDNT) of the Montreal Heart Institute in accordance with the Declaration of Helsinki. Data collection was conducted at the Montreal Heart Institute.

**Table. 1.** Demographic data on participants with gas data.

|  | Controls (37) | CAD (35) | P-value |
| --- | --- | --- | --- |
| Age (years) | 65.35 ± 8.31 | 66.42 ± 9.29 | 0.83 |
| Sex | Female (10) | Female (6) | - |
| Education (years) | 16.22 ± 2.45 | 16.03 ± 4.03 | 0.80 |
| MoCA | 26.73 ± 2.01 | 25.33 ± 3.04 | **0.02** |
| MMSE | 28.25 ± 1.48 | 28.42 ± 1.17 | 0.97 |
| Executive Function(z-score) | 0.314 ± 0.62 | -0.32 ± 1.01 | **<0.01** |
| Working Memory (z-score) | 0.04 ± 0.83 | 0.033 ± 0.79 | 0.95 |
| Processing Speed (z-score) | 0.26 ± 0.72 | -0.34 ± 0.86 | **<0.01** |
| Verbal/Episodic Memory (z-score) | 0.19 ± 0.96 | 0.041 ± 0.97 | 0.31 |
| SBP (mmHg) | 127.12 ± 13.85 | 126.96 ± 14.54 | 0.96 |
| BMI (kg/m<sup>2</sup>) | 26.11 ± 3.94 | 26.95 ± 3.57 | 0.3624 |

\* SBP: Systolic Blood Pressure
\* BMI: Body Mass index
\*MMSE: Mini-Mental State Examination; a brief 30-point test to assess global cognitive function.
\*MoCA: Montreal Cognitive Assessment; a screening tool for mild cognitive impairment covering multiple cognitive domains.

Inclusion criteria for CAD patients were documented coronary artery disease, such as prior acute coronary syndrome, previous coronary angiography or revascularization, or myocardial ischemia documented on myocardial scintigraphy (Table 2 for clinical characteristics). Healthy controls (HCs) were free of cardiac disease, diabetes, and hypertension. HCs had no history of cardiovascular or neurological disease and no current hypertension (resting BP >140/90 mmHg). They were allowed a maximum of two cardiovascular risk factors, excluding age. Individuals with Type I or II diabetes or current hypertension were excluded. All participants had to be fluent in either English or French to ensure informed consent and accurate cognitive assessment.

**Table. 2.** Clinical information in CAD and HC participants.

| **Clinical Health characteristics in patients with CAD (n=35), Healthy Controls (n=37)** |  |  |
| --- | --- | --- |
| **Cardiometabolic risk factors , n(%)** |  |  |
|  | Patients with CAD | Healthy Controls |
| Diabetes | 3 (8) | - |
| Hypertension | 18 (51) | - |
| Dyslipidemia | 28 (80) | 3(8) |
| Obesity | 5(14) | 5(11) |
| Familial history of cardiovascular disease | 21 (60) | 6(16) |
| **Cardiovascular history, n(%)** |  |  |
| Previous MI | 14 (40) | - |
| STEMI | 8 (22) | - |
| NSTEMI | 6 (17) | - |
| Percutaneous coronary intervention (PCI) | 22 (62) | - |
| Coronary artery bypass grafting (CABG) | 10 (28) | - |
| Angina | 10 (28) | - |
| **Cardiovascular medication, n (%)** |  |  |
| Beta-Blocker | 17 (48) | - |
| Aspirin | 11 (31) | - |
| Antilipemic agents (e.g., statins) | 26 (74) | 1(2) |
| RAAS inhibitors | 11 (31) | - |
| Calcium channels blockers | 5 (14) | - |

MI = Myocardial Infarction; STEMI = ST-elevation Myocardial Infarction; NSTEMI = Non-ST-
elevation Myocardial Infarction; RAAS = Renin-Angiotensin-Aldosterone System

Exclusion criteria for all participants included a history of stroke, neurological, psychiatric or respiratory disorder, thyroid disease, potential cognitive impairment (Mini Mental State Examination (MMSE) < 25), tobacco use, high alcohol consumption (more than two drinks per day), and contraindications to MRI (e.g., ferromagnetic implants, claustrophobia). Participants were also excluded if they had undergone surgery under general anesthesia within the past 6 months, had a recent acute coronary event (< 3 months), chronic systolic heart failure, resting left ventricular ejection fraction < 40%, symptomatic aortic stenosis, or severe non-revascularizable coronary artery disease, such as left main coronary stenosis. Other exclusion factors included awaiting coronary artery bypass surgery, having an implanted automatic defibrillator or permanent pacemaker, malignant arrhythmias during exercise, arthritis or claudication, severe exercise intolerance, or excessive discomfort due to hypercapnia (> 5 on the ^29^, dyspnea scale).

Data was collected over four visits. The first visit assessed eligibility via medical history questionnaire and global cognition using the MMSE ^30^, and included a 2-minute hypercapnia test to confirm tolerance for respiratory manipulation (2 participants were excluded). The second visit involved cognitive testing. During the third visit, participants performed a maximal effort test to determine VO₂peak (not reported here). The final visit was dedicated to MRI acquisition.

### MRI acquisition

Data were collected using a 3T Skyra MRI system with a 32-channel head coil. The acquisition protocol included structural MRI, dual-echo pCASL for simultaneous acquisition of perfusion and Bold Oxygen Level Dependent (BOLD) signals and a blood magnetization map (M0) for perfusion quantification. pCASL data was acquired with a voxel resolution of 3.4375 x 3.4375 x 7 mm, TR/TE1/TE2/alpha: 4000/10/30ms/90°, labeling duration of 1517 ms with a post-labeling delay (PLD) of 1300 ms. The M0 acquisition had identical parameters but with a TR of 10 seconds, to ensure a fully recovered magnetization.

The MRI session was split into two sub-sessions to allow mask removal during part of the scan. Two T1-weighted images were acquired to ensure accurate pCASL slab registration: a lower-resolution scan before pCASL with the mask on (MPRAGE, TR/TE/Flip angle = 15 ms/3.81 ms/25°, 1.5 mm isotropic), and a higher-resolution scan in the second sub-session without the mask (MPRAGE, TR/TE/Flip angle = 2300 ms/2.32 ms/8°, 0.9 mm isotropic).

### Respiratory manipulation

The RespirActTM system (RespirAct_TM_, Thornhill Research, Toronto, Canada) was used during the breathing manipulation to target specific end-tidal partial pressures of CO₂ and O₂. Three conditions were sequentially targeted for 2 minutes, preceded and followed by 2 minutes of air inhalation: a hypercapnic condition (targeting 5 mmHg above baseline for end-tidal partial pressure CO₂ (PETCO₂)), an isocapnic hyperoxic condition (targeting 150 mmHg for end-tidal O₂), and a combined hyperoxia and hypercapnia condition (targeting 5 mmHg above baseline for PETCO₂ and 150 mmHg for PETO₂). A rebreathing face mask was used to deliver gases to the participants. The system delivered gas at a flow rate of 20 L/min, while the concentrations of exhaled CO_2_ and O_2_ were monitored. On their first study visit, a familiarization session was done with a 2min hypercapnia manipulation to ensure participant comfort during the MRI. Breathing discomfort was assessed using the Banzett dyspnea scale and those with a score >5 were not invited to continue in the study (n = 2)^29^.

### Cognitive assessment

A comprehensive neuropsychological assessment was administered in the following order. Global cognition was first assessed using the Montreal Cognitive Assessment (MoCA). Verbal and auditory episodic memory was then evaluated using the Rey Auditory Verbal Learning Test (RAVLT) ^31^. Working memory was assessed with the Digit Span (DS) tests, forward and backward ^32^. Next, processing speed was measured using the Digit Symbol Substitution Test (DSST) ^33^. Subsequently, the Delis-Kaplan Executive Function System (D-KEFS) Color Word Interference Test (CWIT) ^34^, consisting of four conditions (color naming, reading, inhibition, and switching), was conducted. Finally, the Trail Making Test (TMT) ^35^, comprising part A (processing speed) and part B (executive functioning), was administered, requiring participants to connect numbers sequentially (part A) and alternate between numbers and letters sequentially (part B).

All raw test scores were transformed into standardized z-scores, from which four cognitive domain composite scores were created, as in ^36^. The executive functioning composite score was computed as the mean of the TMT Part B and CWIT inhibition and switching conditions z-scores. The processing speed composite score was derived from the mean of the DSST (multiplied by -1), TMT Part A, and the CWIT color naming and reading conditions z-scores. The working memory composite was calculated as the mean of the z-scores of the DS forward and backward tests, while the verbal/episodic memory composite was determined by averaging the z-scores from the RAVLT total five learning trials, immediate recall, and delayed recall. To facilitate interpretation, each score derived from response speed was multiplied by -1 so that higher scores indicate better cognitive performance. Internal consistency for each cognitive composite was evaluated using Cronbach’s alpha. Reliability was excellent for Verbal Memory (α = 0.94), good for Executive Function (α = 0.80) and Processing Speed (α = 0.79), but lower for Working Memory (α = 0.50), indicating limited internal consistency within that particular domain.

### Respiratory data analysis

CO_2_ and O_2_ were sampled continuously throughout the breathing manipulation by the RespirAct, which also generates a time series of end-tidal CO_2_ and O_2_ partial pressures. The end-tidal partial pressure is used as a proxy for arterial gas partial pressure ^37^. To combine these values with the MRI data, a MATLAB script was employed to smooth the data, remove outliers and resize the data to match the durations of the BOLD and ASL signals.

### Imaging data processing

### T1 processing

All structural images underwent preprocessing with the Brain Extraction Tool (BET) in FSL to remove the skull. Subsequently, FSL’s FAST ^38^ was employed to segment the structural images into Grey Matter (GM), White Matter (WM), and Cerebrospinal Fluid (CSF).

### Preprocessing of BOLD and pCASL

Both ASL and BOLD datasets were motion-corrected using FSL’s mcflirt. A brain mask was created from the motion-corrected BOLD images using BET and applied to both echoes to remove the scalp. The first echo of the dual-echo pCASL acquisition was used to compute the perfusion-weighted time-series, from which CBF was estimated via surround subtraction. Volumes with voxel intensities exceeding 3 standard deviations from the mean were flagged, and those with >50% outlier voxels were excluded to ensure sufficient tSNR ^39^. The M0 image was skull-stripped using BET and used for perfusion calibration. The second echo was used to extract the BOLD-weighted signal via surround addition, reflecting changes in blood oxygenation.

### CBF and CVR maps

Perfusion was quantified from the preprocessed ASL time-series using FSL’s BASIL with kinetic modeling and M0 acquisition. Partial volume correction was applied via weighted averaging based on tissue contributions from GM, WM, and CSF segmentations of the T1-weighted data ^40^. CVR was quantified from the BOLD time-series derived from the second echo of the pCASL data using surround addition. Due to dead space in the tubing, there is an individual-specific delay between gas sampling and the corresponding brain MRI signal, requiring adjustment of end-tidal traces. This lag was estimated via cross-correlation using a custom MATLAB script, selecting the delay where PETCO_2_ and the mean GM BOLD signal were maximally correlated ^41^. Absolute CVR was calculated as the percent BOLD signal change per mmHg CO_2_ from a General Linear Model (GLM) using the ETCO_2_ time course as the regressor.

### OEF & CMRO_2_ maps

OEF was estimated using the general calibration model (GCM), based on the deoxyhemoglobin dilution model of the BOLD signal ^42^. Detailed equations and the full estimation process for OEF are provided in the Supplementary material. Following OEF estimation, CMRO_2_ was calculated using CBF maps and arterial oxygen content using Fick’s principle, as described in Equation 7 of the supplementary material.

### Registration to MNI space

The pCASL images were registered to MNI space in three separate steps. First, a rigid body registration with nearest neighbor interpolation was conducted using Advanced Normalization Tools (ANTs) to align the pCASL mean images with the low-resolution (1.5 mm) T1-weighted image. Next, the low-resolution T1-weighted images were registered to the high-resolution (0.9 mm) T1 space using a multistage approach that combined rigid, affine, and nonlinear (SyN) transformations with a multi-resolution strategy. In the final step, the high-resolution T1 images were registered to MNI space using the same transformation method applied for the registration from low to high resolution. Finally, the pCASL images were registered to MNI space by applying the three transformation matrices obtained from the previous registration steps, utilizing 3 warp images.

### Statistical Analysis

### Patients vs controls

We performed a region of interest (ROI)-based analysis to identify brain regions most affected by CAD, using the LPBA40 atlas comprising 56 regions across frontal, temporal, parietal, and subcortical areas ^43^. Due to low tSNR in the occipital lobe ASL data, occipital, lingual, and fusiform regions were excluded, yielding 44 regions plus total gray matter (45 in total). Outliers were defined as values exceeding three standard deviations from the mean. Linear regressions in R assessed group differences in mean regional values, controlling for age and sex. False Discovery Rate (FDR) correction was applied, with significance set at p < 0.05.

### Correlation Between Cognitive Composite Scores, and Brain Biomarkers

Linear regressions in R were used to investigate the relationships between cognitive composite scores and regional CBF, CVR, OEF and CMRO_2_. Age, sex, group (CAD and control) and education were included as covariates. All p-values were adjusted for multiple comparisons using FDR with a significance threshold of p < 0.05.

## Results

### Patients vs Controls

Table 1 shows the demographic for each group. Group comparisons revealed no significant differences in age, education, BMI, or systolic blood pressure between the CAD and control groups. However, participants with CAD demonstrated significantly lower global cognitive function as assessed by the MoCA (p = 0.02), executive function (p < 0.01) and processing speed (p < 0.01). No group differences were observed in MMSE scores, working memory, or verbal episodic memory performance. Clinical profiles for the CAD and healthy control groups are summarized in Table 2.

### Cerebrovascular health

CAD patients exhibited reduced CBF across the whole GM (p = 0.036) and in multiple brain regions both in cortical and subcortical regions (see Table 3 and Figure 1). CVR was also significantly lower in patients across the GM (p < 0.01), with local reductions in several cortical regions (see Table 3 and Figure 1).

**Figure. 1.**
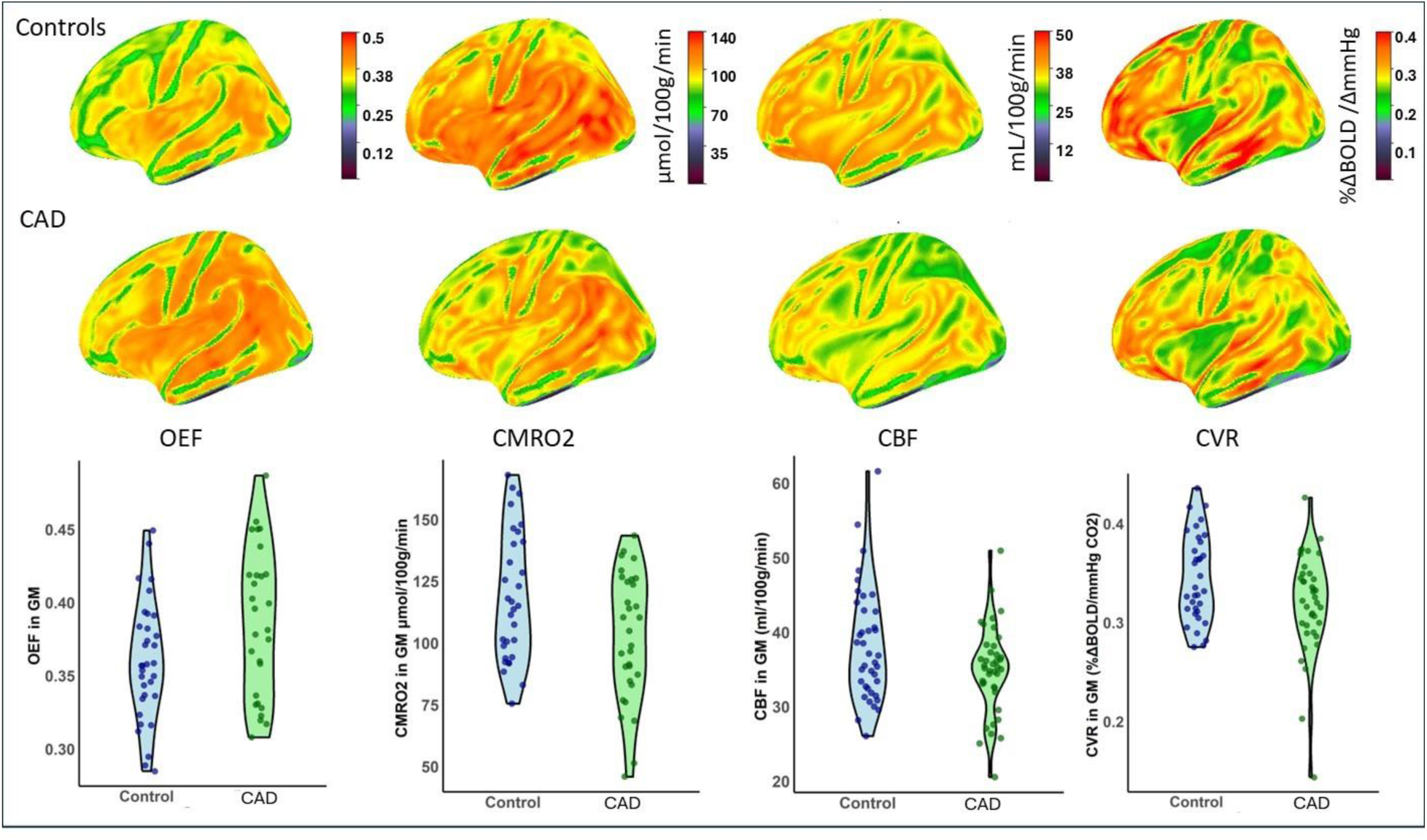
Surface maps showing CBF, CVR, CMRO2, and OEF in both patients and controls, along with corresponding violin plots for the whole GM average for each group.

**Table 3.** Brain regions with significant differences in CBF (mL/100g/min) and CVR (%ΔBOLD/ΔmmHg).

|  | CBF |  |  | CVR |  |  |
| --- | --- | --- | --- | --- | --- | --- |
|  | Controls | CAD | P<br>value | Controls | CAD | P<br>value |
| GM | 38.2 ± 7.6 | 34.9 ± 5.85 | **0.03** | 0.34 ± 0.04 | 0.31 ± 0.03 | **<0.01** |
| L Sup Frontal G | 34.3 ± 6.22 | 30.8 ± 5.35 | **0.01** | 0.32 ± 0.04 | 0.28 ± 0.04 | **<0.01** |
| R Sup Frontal G | 33.9 ± 5.9 | 30.9 ± 5.25 | **0.02** | 0.31 ± 0.04 | 0.28 ± 0.04 | **<0.01** |
| L Mid Frontal G | 34.1 ± 6.81 | 31.8 ± 5.62 | n. s | 0.29 ± 0.05 | 0.27 ± 0.04 | **0.02** |
| R Precentral | 34.5 ± 7.02 | 31.6 ± 5.45 | **0.04** | 0.27 ± 0.05 | 0.26 ± 0.05 | n. s |
| L Precentral | 34.0 ± 6.37 | 31.6 ± 5.28 | n. s | 0.28 ± 0.04 | 0.26 ± 0.04 | **0.04** |
| L Mid Orbital G | 37.8 ± 7.29 | 32.8 ± 6.74 | **<0.01** | 0.33 ± 0.05 | 0.31 ± 0.06 | n. s |
| R Mid Orbital | 36.9 ± 6.03 | 33.4 ± 6.55 | **0.02** | 0.32 ± 0.06 | 0.31 ± 0.05 | n. s |
| L Lat Orbital G | 38.6 ± 6.03 | 33.4 ± 6.55 | **0.01** | 0.34 ± 0.05 | 0.31 ± 0.06 | **0.02** |
| R postcentral G | 34.5 ± 7.33 | 31.0 ± 5.33 | **0.02** | 0.26 ± 0.04 | 0.26 ± 0.05 | n. s |
| R precuneus | 34.5 ± 7.02 | 31.6 ± 5.45 | n. s | 0.37 ± 0.05 | 0.34 ± 0.05 | **0.02** |
| L Sup Temporal G | 37.3 ± 8.05 | 33.8 ± 6.20 | **0.04** | 0.33 ± 0.05 | 0.30 ± 0.04 | **0.04** |
| L Insula | 38.5 ± 9.92 | 33.7 ± 6.81 | **0.01** | 0.27 ± 0.06 | 0.26 ± 0.06 | n. s |
| R Insula | 37.6 ± 8.07 | 33.8 ± 6.75 | **0.03** | 0.26 ± 0.06 | 0.25 ± 0.06 | n. s |
| L Cingulate G | 33.5 ± 7.56 | 30.2 ± 6.07 | **0.04** | 0.28 ± 0.05 | 0.27 ± 0.06 | n. s |
| L Caudate | 29.4 ± 6.43 | 26.0 ± 4.13 | **<0.01** | 0.27 ± 0.06 | 0.26 ± 0.07 | n. s |
| L Putamen | 31.8 ± 6.04 | 35.6 ± 7.97 | **0.02** | 0.24 ± 0.05 | 0.24 ± 0.06 | n. s |
| R Putamen | 34.05 ± 7.48 | 30.5 ± 5.75 | **0.03** | 0.23 ± 0.05 | 0.24 ± 0.05 | n. s |

\* Regions with significant differences in both CBF and CVR are highlighted in bold adjusted by age and sex as covariates.
\* GM: Grey Matter; L: left ; Sup: superior; Mid: Middle; G: gyrus; Lat: lateral

### Cerebral metabolic health

We observed significantly lower CMRO₂ in CAD patients across the whole GM (p = 0.01), as well as in several cortical and some subcortical regions (see Table 4 and Figure 1). OEF was significantly higher in the CAD group compared to controls in the whole GM (p = 0.02) and in multiple cortical areas(Table 4 and Figure 1). The spatial distribution of significant regions for all biomarkers is presented in Figure S1.

**Table. 4.** Brain regions showing significantly higher OEF and lower CMRO_2_ (μmol/100g/min) in patients.

|  | OEF |  |  | CMRO<sub>2</sub> |  |  |
| --- | --- | --- | --- | --- | --- | --- |
|  | Controls | CAD | P value | Controls | CAD | P value |
| GM | $0.36 \pm 0.04$ | $0.38 \pm 0.05$ | **0.02** | $118.4 \pm 25.8$ | $102.6 \pm 25.6$ | **0.01** |
| R Mid Orbital G | $0.32 \pm 0.06$ | $0.32 \pm 0.06$ | n. s | $109.9 \pm 25.8$ | $93.07 \pm 29.9$ | **0.02** |
| R Lateral Orbital G | $0.32 \pm 0.06$ | $0.33 \pm 0.04$ | n. s | $113.8 \pm 30.3$ | $96.9 \pm 28.3$ | **0.01** |
| L precuneus | $0.37 \pm 0.05$ | $0.38 \pm 0.08$ | n. s | $117.5 \pm 31.1$ | $99.7 \pm 30.2$ | **0.02** |
| R precuneus | $0.36 \pm 0.05$ | $0.37 \pm 0.08$ | n. s | $117.7 \pm 32.3$ | $99.2 \pm 30.9$ | **0.01** |
| L precentral G | $0.31 \pm 0.06$ | $0.36 \pm 0.05$ | **<0.01** | $98.3 \pm 27.0$ | $94.6 \pm 25.8$ | n. s |
| R precentral G | $0.32 \pm 0.06$ | $0.36 \pm 0.05$ | **0.02** | $104.9 \pm 27.7$ | $95.3 \pm 26.08$ | n. s |
| R postcentral G | $0.33 \pm 0.06$ | $0.37 \pm 0.05$ | **0.02** | $107.5 \pm 27.7$ | $95.3 \pm 26.2$ | n. s |
| L Sup Temporal G | $0.32 \pm 0.05$ | $0.36 \pm 0.05$ | **<0.01** | $113.2 \pm 26.8$ | $102.8 \pm 28.1$ | n. s |
| R Sup Temporal G | $0.34 \pm 0.05$ | $0.36 \pm 0.06$ | n. s | $117.9 \pm 26.5$ | $102.6 \pm 28.4$ | **0.03** |
| L Sup Parietal G | $0.32 \pm 0.05$ | $0.36 \pm 0.06$ | **0.01** | $97.6 \pm 26.3$ | $88.4 \pm 23.1$ | n. s |
| L Sup Marginal G | $0.33 \pm 0.05$ | $0.37 \pm 0.07$ | **0.01** | $111.5 \pm 30.4$ | $106.8 \pm 31.3$ | n. s |
| L Angular G | $0.33 \pm 0.05$ | $0.38 \pm 0.07$ | **<0.01** | $106.2 \pm 30.1$ | $105.0 \pm 34.3$ | n. s |
| R Cingulate G | $0.38 \pm 0.05$ | $0.38 \pm 0.07$ | n. s | $112.9 \pm 30.52$ | $92.3 \pm 27.3$ | **<0.01** |
| L Cingulate G | $0.38 \pm 0.05$ | $0.38 \pm 0.06$ | n. s | $110.5 \pm 32.3$ | $89.3 \pm 26.4$ | **<0.01** |
| R Insula | $0.38 \pm 0.07$ | $0.39 \pm 0.07$ | n. s | $121.8 \pm 34.1$ | $100.6 \pm 34.4$ | **0.02** |

\*Significance was defined as $p < 0.05$ after FDR correction. Age and sex were included as the covariates in the analysis.
\* CAD: Coronary Artery disease group
\* GM: Grey Matter; L: left ; Sup: superior; Mid: Middle; G: gyrus.

### Relationship with cognition

We then explored the relationships between measures of cerebral vascular and metabolic health and cognitive performance across four domains: executive function, working memory, processing speed, and episodic memory. No significant associations were found between cognitive performance in any domain and CMRO_2_ or CBF. However, we identified several regions where CVR was positively correlated with executive function (Figure 2 and Table 5), particularly in the whole GM and in frontal, temporal, parietal and a few subcortical regions such as the hippocampus and putamen. Additionally, we observed a significant negative relationship between OEF in frontal and temporal regions and working memory (Figure 3 and Table 5).

**Figure 2.**
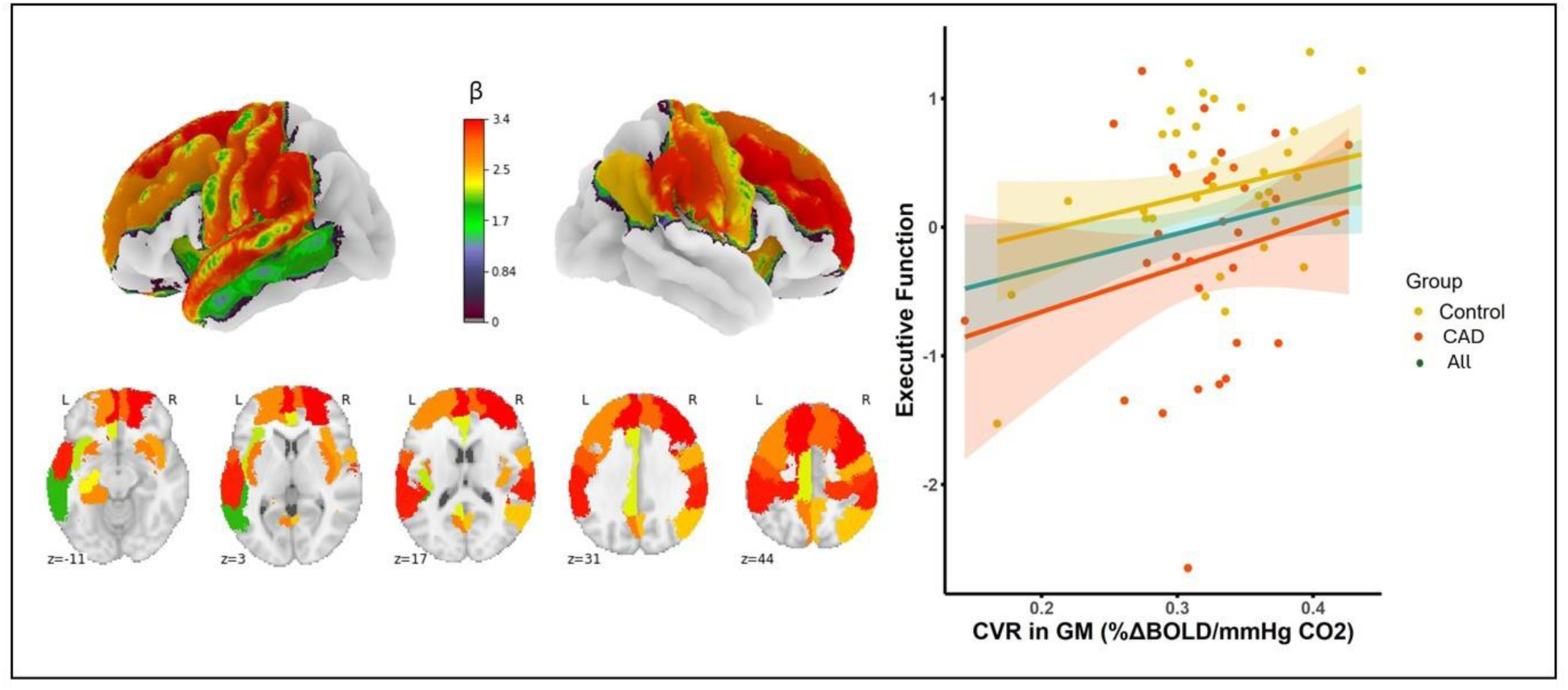
Brain surface visualization highlighting regions with significant relationships between CVR and executive function. The color gradient represents beta values from the linear regression analysis for all participants. Additionally, linear plots illustrate the relationship between executive function and CVR within GM, presented for all participants combined (green line), as well as for each group separately (yellow for controls and red for CAD).

**Figure 3.**
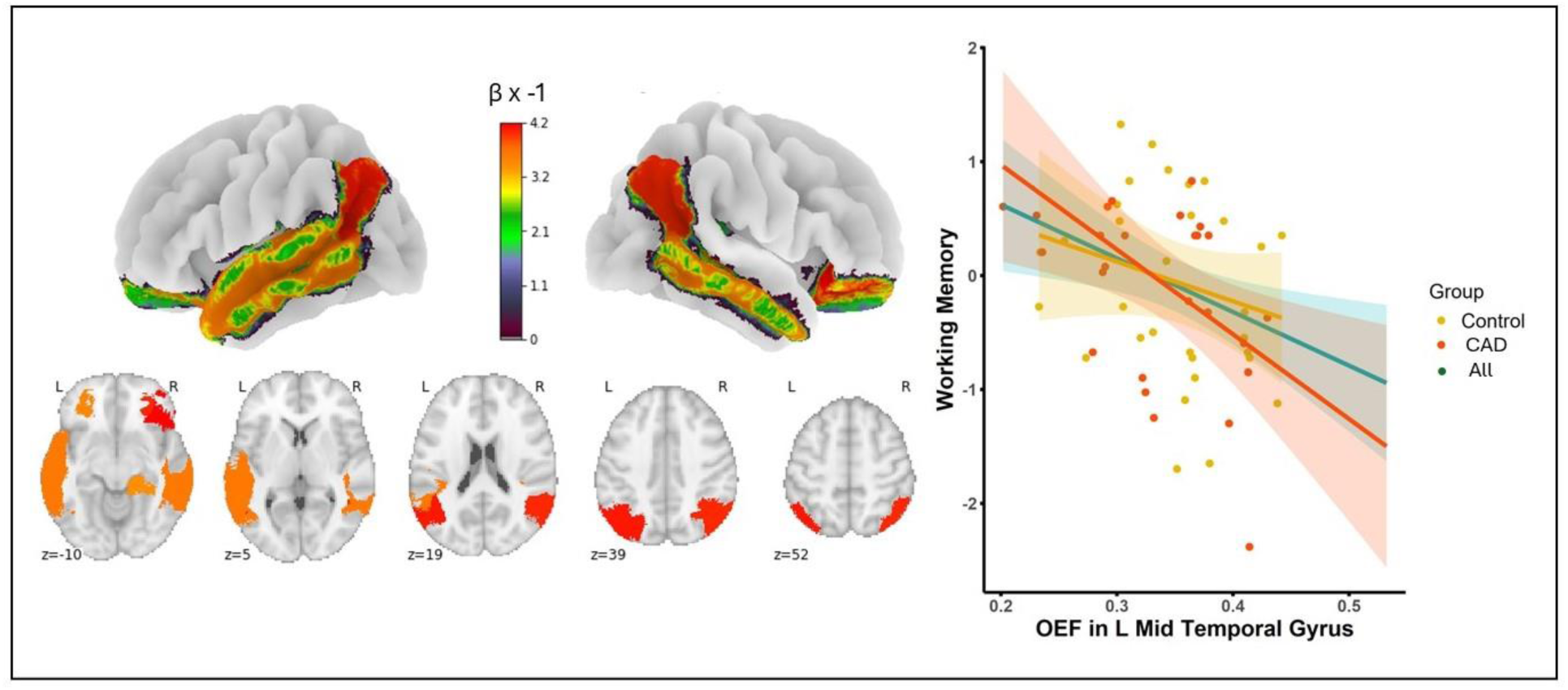
Brain surface visualization highlighting regions with significant relationships between OEF and working memory. The color gradient represents beta values (multiplied by -1 so that all values are positive for visualization) from the linear regression analysis for all participants. Additionally, linear plots illustrate the relationship between working memory and OEF in the medial temporal gyrus, presented for all participants combined (green line), as well as for each group separately (yellow for controls and red for CAD).

**Table. 5.**
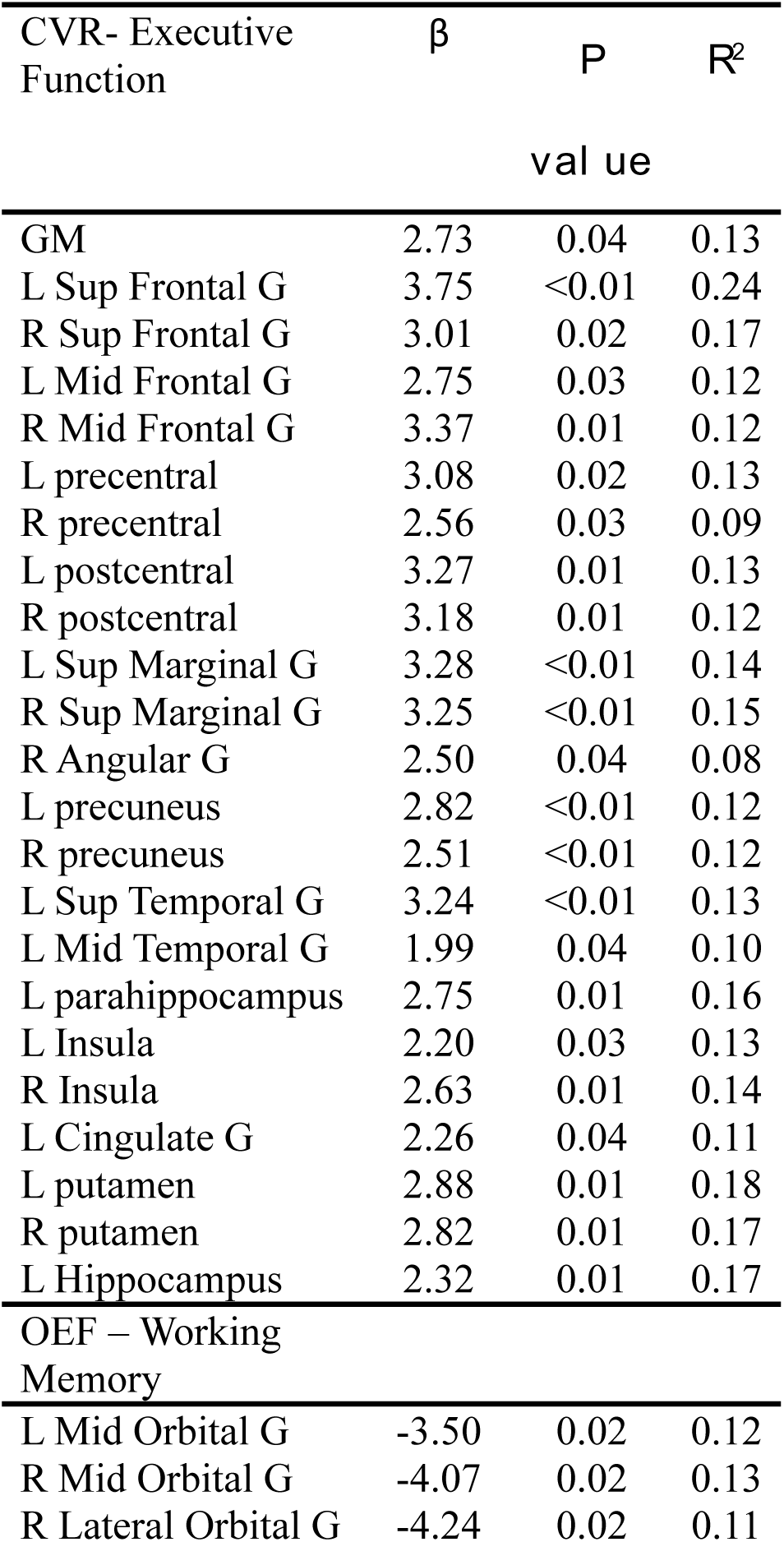
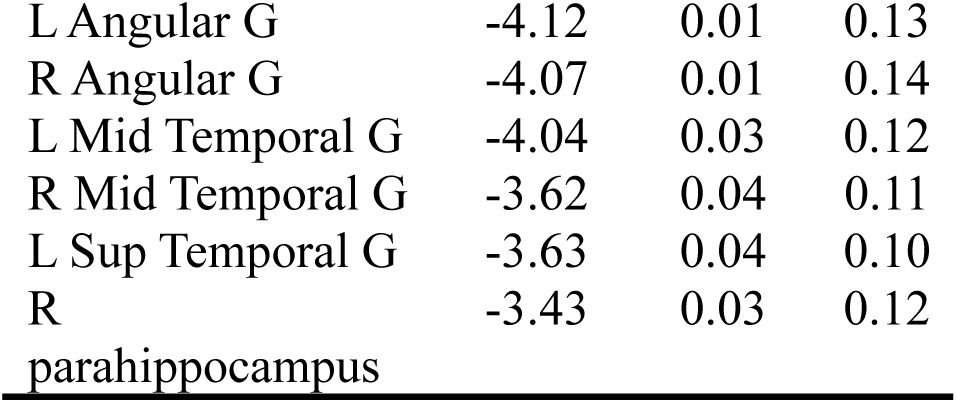
Brain regions showing significant correlations between CVR and executive function in the top half of the table and between OEF and working memory in the bottom half with age, sex, group and education as covariates.

Finally, in all analyses linking brain outcomes and cognition, group interactions (controls vs CAD) were all nonsignificant, indicating that these relationships may not be specific to one group.

## Discussion

Our findings reveal that in our sample, CAD is associated with profound and widespread impairments in both cerebral vascular and metabolic health, with patients showing lower CBF, CVR, and CMRO_2_, and higher OEF as compared to healthy controls. In our sample, CAD was also associated with lower global cognition (MoCA), as well as reduced performance in executive functions and processing speed. We uncovered relationships between brain physiology and cognition, showing a positive relationship between executive function and CVR, and a negative association between working memory and OEF in the whole group. Importantly, these associations between CVR, OEF and cognition are not unique to our CAD patients, but are also present in healthy individuals. These results represent the most comprehensive portrait of the impact of CAD on brain physiology to date, and highlight the role of vascular reserve (i.e. CVR) and the balance between oxygen consumption and delivery (i.e. OEF) in maintaining cognitive function.

### Patients vs controls

Reduced CBF and CVR in our CAD patients are consistent with the few other studies that have investigated cerebrovascular health in CAD. Anazodo et al. ^10^ observed localized CBF deficits, within the larger regions identified in our sample. Our results therefore show that more widespread deficits exist in this population than previously thought, likely due to the greater power of our larger dataset. In contrast, the CVR impairments identified here were in distinct regions, perhaps due to the different methods used in those studies (ASL-based CVR in Anazodo et al., and BOLD-based CVR here). The higher signal-to-noise ratio of BOLD, in addition to the larger sample size, may partly explain the more widespread deficits identified in our data. CVR reductions had also been identified by Haratz et al. ^13^ in the MCA territory, in agreement with the location of several regions showing reduced CVR here ^44^, though our whole brain approach allowed us to identify additional brain regions.

These deficits occur in the larger context of normal aging-related CBF and CVR declines ^45–47^, and could arise from several systemic and cerebrovascular factors. CAD is associated with both atherosclerosis and endothelial dysfunction ^48^. Prior studies suggest that atherosclerosis and endothelial dysfunction could result in reduced nitric oxide availability, the main pathway for CO_2_-mediated vasodilation ^49^, higher vascular stiffness (which could elevate cerebrovascular resistance), leading to impairments in cerebral perfusion and vasodilation ^18,50–52^. Additionally, in cases where cardiac output is reduced—a common, though not universal condition in CAD— cerebral blood flow could be further compromised ^53,54^. Consequently, it is plausible that these vascular alterations are present to various extent in our CAD group and negatively affect both CBF and CVR. However, given that these mechanisms were not directly measured in our study, further research is necessary to explore the mechanisms underlying the cerebrovascular changes observed.

In contrast to CBF and CVR, cerebral metabolic alterations in CAD remain largely unexplored, and to our knowledge, this is the first study to investigate the impact of CAD on regional OEF and CMRO₂. Here, we document an elevated OEF and lower CMRO_2_ in individuals with CAD as compared to age-matched controls, in whole grey matter, with frontal, parietal and temporal regions being most affected. This result is consistent with reduced metabolism, accompanied by a reduced or less efficient blood supply (i.e. reduced CBF), which for a given CMRO_2_ leads to a higher fraction of O_2_ being extracted from the blood to attempt to meet the metabolic demand (i.e. a higher OEF). This result is consistent with an additional metabolic deficit beyond the more modest decline observed across several studies with healthy older adults ^25,55^.

The case of OEF is more complex. We observed a higher OEF in CAD, consistent with a reduced blood supply for a given O_2_ consumption, potentially indicating tissue at risk for ischemic damage ^28^. This elevated OEF is consistent with results by McFadden et al. (2025)^27^, showing higher OEF in individuals with a high cardiometabolic risk factor load, though in their case no concomitant CMRO_2_ reduction was observed. Because their cohort had less advanced vascular pathology, this may indicate that in preclinical or very early vascular compromise, OEF compensation may be sufficient to preserve oxidative metabolism. The elevated OEF observed here is also consistent with the higher OEF observed in Alzheimer’s patients with higher cardiometabolic risk factor load ^56^, and the elevated OEF observed in older adults as compared to younger adults ^25,57^, especially in males ^58^.

Because we concurrently observe a higher OEF and lower CMRO_2_ in CAD patients, these results are consistent with two possible scenarios. In the first, inadequate perfusion has led to cell death and therefore reduced metabolic demand. In the second scenario, while blood supply is reduced and OEF increased, there is additionally reduced mitochondrial efficiency which decreases CMRO_2_ without tissue loss. Substrates are provided, but mitochondria are unable to use the oxygen provided to power the cell, which either has reduced metabolism overall, or a greater contribution of non-oxidative metabolic pathways. In either case, our results are consistent with a confluence of both cerebral vascular and metabolic impairments in CAD. Future studies in larger samples with a wider range of vascular risk and disease which also document mitochondrial dysfunction would help establish the mechanisms that underlie the deficits in OEF and CMRO_2_ we observed in our CAD patients.

### Links with cognition

Despite CBF and CMRO_2_ having the most widespread deficits in CAD in our sample, CBF and CMRO_2_ were not related to any cognitive outcomes. Although this aligns with some previous research ^27,59,60^, it is inconsistent with studies that have reported a positive relationship between CBF and cognitive function in healthy participants ^61^, Alzheimer’s disease ^17^ and vascular cognitive impairment ^19^. In contrast, the positive relationship between CVR and executive function and the negative association observed between OEF and working memory indicate that CVR and OEF may be more sensitive biomarkers of functionally impaired brain health.

Our results showing reduced executive function performance in CAD (in our sample) is consistent with the literature, showing frequent deficits in this domain ^6^. Our results suggest that cerebrovascular deficits, especially in CVR, may underlie this poorer executive function performance. This link between reduced CVR with poorer executive function performance is consistent with one prior study in CAD ^13^. It is also in line with emerging evidence positioning CVR as an early and sensitive biomarker of cognitive impairment in Alzheimer disease and cerebral small vessel disease ^62–64^. Our findings suggest that this relationship between CVR and executive function is present regardless of cardiovascular status. While both healthy controls and our CAD patients show a similar positive relationship between CVR and executive performance, our patients with CAD tend to operate from a lower CVR baseline. This reduced vascular reserve may leave them more susceptible to cognitive decline, positioning them effectively further along within the same trajectory of cerebrovascular impairment-related cognitive deficits. Importantly, our results highlight the promise of CVR as a brain-specific biomarker to be used in future interventions targeting improvements in executive functions.

Our novel finding of a negative association between OEF and working memory offers new insight into the impact of vascular/metabolic compromise on cognitive function in CAD patients. While a recent study involving individuals with cardiovascular risk factors reported no association between OEF and global cognition (as measured by MoCA)^27^, this may be attributable to their smaller sample size and the absence of domain-specific cognitive assessments. Notably, although prior research demonstrated a link between temporal CBF and working memory deficits in healthy participants and people with vascular cognitive impairment ^19,65^, our findings are the first to reveal a similar pattern for OEF. Notably, we did not observe any relationship between CBF and working memory in our data, indicating that CBF per se is not as crucial as a mismatch between perfusion and the underlying oxidative demand. This is consistent with elevated OEF being a better marker of ischemic risk, given that it captures aspects of both demand and delivery.

In our sample, CAD patients exhibited lower resting-state CBF and CMRO₂, but these baseline reductions were not associated with cognitive performance. Instead, cognitive scores were more closely related to CVR and OEF, which are more related to the brain’s capacity to respond to metabolic challenges. CVR reflects the cerebrovascular reserve, or the capacity of vessels to dilate in response to increased metabolic demand, whereas OEF indicates the extent to which tissue already maximizes oxygen extraction from available blood flow, resulting in low mitochondrial oxygen tension. If cerebrovascular reserve is diminished or if oxygen extraction is already nearing maximal capacity, the brain’s ability to accommodate transient increases in metabolic demand may become limited, resulting in reduced cognitive performance. This interpretation aligns with previous studies emphasizing the importance of cerebrovascular adaptability in supporting cognitive resilience during aging and disease ^18,59,60^.

## Limitations and future directions

This study provides the most comprehensive characterization to date of brain physiological deficits linked to CAD and their relationship with cognitive function. However, several limitations should be noted. First, the lack of cardiac function quantification limits insights with regards to mechanistics involved in the interplay between cardiac performance and brain health. Future studies should incorporate direct measures of cardiac function to better disentangle heart-and brain-specific contributions. Second, the sample was overly homogeneous—recruited from a single site, predominantly male, and excluding those with severe CAD—limiting generalizability. Broader, more diverse cohorts are needed to validate these findings. Third, ASL inherently has low tSNR, further affected by prolonged arterial transit times in vascular-risk populations. The use of a single PLD may underestimate CBF in individuals with delayed transit. Future studies should employ multi-PLD sequences and background suppression to enhance CBF accuracy. While background suppression was omitted here to preserve BOLD SNR for CVR, OEF, and CMRO₂ estimation, future work may use dual excitation methods to allow its inclusion.

## Conclusion

Our findings show that CAD in our sample is associated with widespread impairments in cerebral vascular and metabolic health, characterized by lower CVR, CBF and CMRO_2_, and elevated OEF. Poorer CVR and OEF were linked to worse executive function and working memory, suggesting that maintenance of vascular reserve and the balance between oxygen usage and delivery is crucial for preserving cognition. This research highlights the need to target cerebral vascular and metabolic health in preventative interventions to maintain cognition in CAD.

## Acknowledgments

We would like to thank everyone who contributed to this project: Paule Samson, Thomas Vincent, Julie Lalongé, Hakima Benhalima, Milla Shakleva, Victoria D’Amours, Agathe Godet, Stephanie Beram, Roni Zaks, Robert Hovey, Alexandre Bailey, Catherina Medeiros, and Zineb Rouabah. Thank you also to the laboratories of Dr Louis Bherer and Dr Mathieu Gayda. Lastly, we would like to acknowledge our research participants without whom none of this would have been possible.

## Supplemental Material

Supplemental Methods

Table S1

Figure S1

## Nonstandard Abbreviations and Acronyms

CAD: coronary artery disease
MRI: magnetic resonance imaging
CBF: cerebral blood flow
CVR: cerebrovascular reactivity
OEF: oxygen extraction fraction
CMRO_2_: cerebral metabolic rate of oxygen
HC: healthy controls
PETCO_₂_: end-tidal partial pressure of CO₂
PETO_₂_: end-tidal partial pressure of O₂

